# Distinct encoding and post-encoding representational formats contribute to episodic sequence memory formation

**DOI:** 10.1101/2022.08.09.503295

**Authors:** Xiongbo Wu, Lluís Fuentemilla

**Affiliations:** Department of Cognition, Development and Educational Psychology, University of Barcelona, Barcelona, Spain; Institute of Neurosciences, University of Barcelona, Barcelona, Spain; Department of Psychology, Ludwig-Maximilians-Universität München, Munich, Germany; Cognition and Brain Plasticity Unit, Bellvitge Institute for Biomedical Research, Hospitalet de Llobregat, Spain

## Abstract

In episodic encoding, an unfolding experience is rapidly transformed into a memory representation that binds separate episodic elements into a memory form to be later recollected. However, it is unclear how brain activity changes over time to accommodate the encoding of incoming information. This study aimed to investigate the dynamics of the representational format that contributed to memory formation of sequential episodes. We combined Representational Similarity Analysis and multivariate decoding approaches on EEG data to compare whether “category-level” or “item-level” representations supported memory formation during the online encoding of a picture triplet sequence and offline, in the period that immediately followed encoding. The findings revealed a gradual integration of category-level representation during the online encoding of the picture sequence and a rapid item-based neural reactivation of the encoded sequence at the episodic offset. However, we found that only memory reinstatement at episodic offset was associated with successful memory retrieval from long term memory. These results suggest that post-encoding memory reinstatement is crucial for the rapid formation of unique memory for episodes that unfold over time. Overall, the study sheds light on the dynamics of representational format changes that take place during the formation of episodic memories.

## Introduction

An important challenge in neuroscience is to understand how learning systems operate to rapidly transform an ongoing experience into bound episodic memory traces that can be recollected at long term.

Traditionally, human episodic memory research answered this question by focusing on experiments using single items as studied material and analysing the neural underpinnings that predicted their successful retrieval during the online encoding (i.e., when stimuli to be remembered were present) (Paller & Wagner, 2002). This research showed that the same brain regions and patterns of activity that are engaged during memory “encoding” of an item tend to be reinstated during subsequent memory “retrieval” (Danker et al., 2017; Gordon et al., 2014; Ritchey et al., 2013; Staresina et al., 2012, 2016), suggesting that remembering relies on reactivating the initial neural representations elicited online during encoding. Alternatively, another set of studies using single pictures as a studied material has emphasized that successful memory encoding involves a substantial transformation of early representations elicited during perception (Liu et al., 2021). Indeed, it has been shown that within the first few hundred milliseconds, brain activities gradually and progressively change from representing low-level visual information to higher-order categorical and semantic information (Clarke et al., 2018) and that these transformed, semantic representational formats contributed to stable short-term memory maintenance (Liu et al., 2020). While these two lines of research evidence highlight the dynamic representational nature that accounts for how the brain encodes single items in memory, it is also relevant to understand how these representational dynamics operate in the context of a sequential episodic encoding, akin to a more realistic scenario whereby the retrieval of an event involves remembering the elements that were encoded in the unfolding experience.

In the current study, we sought to investigate the representational format that accounts for successful encoding of episodic sequences in memory. The use of episodic sequences allows us to investigate an important question that has been largely overlooked in the literature: how does brain activity change over time to accommodate the encoding of incoming information? Specifically, we aimed to examine whether successful encoding is supported by the preservation of the representational acuity or idiosyncrasy of each of the elements within a sequence or, alternatively, if it better requires an undergoing representational transformation of each of the elements so that they become integrated into a bound but semantic representational structure. In addition, we also aimed to investigate whether the immediate offline period after the completion of a sequential episode contributes to successful encoding. This is motivated by recent research that showed that brain mechanisms that followed online encoding of a continuous stream of stimuli offer an “optimal” window to register in memory a bound representation of an episode (e.g., Lu et al., 2022). Here, we hypothesize that one such offline neural mechanism promoting the binding of the elements of a sequence episode in memory is the rapid reactivation of them in a temporally compressed manner. Rodent studies showed that neural replay at quiescent periods immediately after single-trial spatial experiences supports the formation of high-fidelity representation of the encoded trajectory in memory (Foster & Wilson, 2006). More recently, research in rodents pointed to a semantic aspect to replay, since it has been shown that novel information is added to the replay memory of a track upon subsequent re-exposures to it (Berners-Lee et al., 2022). In line with the animal literature, human literature has recently shown that representations of just-ended events are reactivated upon the completion of an event (Wu et al., 2022) or at event boundaries (Silva et al., 2019; Sols et al., 2017). However, these studies did not discern whether memory reinstatement of the just-encoded episode and its impact on retrieval relies on reactivating the initial neural representations elicited online during encoding or, alternatively, whether it strengthens the transformation of early encoded episodic material into a more semantic, abstract representation. Thus, an important question that remains unresolved relates to the representational format of this offset-locked neural activity supporting later recollection of the just-encoded experience.

We recorded scalp electrophysiological (EEG) signals while participants encoded trial-unique combinations of face-object-scene picture triplet sequences to be subsequently recalled in a test. Leveraged by the high temporal resolution of the EEG recording and the analytical power of representational similarity analysis (RSA) and multivariate decoding, we examined the dynamics of the representational format that contributed to successful episodic memory formation. The combination of these two analytical approaches allowed us to discern whether different types of memory representations, namely, “category-level” and “item-level”, supported memory during the online encoding of the triplet sequence and offline, in the period that immediately followed encoding. Our results demonstrate the time course, and the functional relevance of the different representational formats that have an impact on memory formation and help advance our understanding of how the brain rapidly transforms the unfolding experience into bound episodic memory traces.

## Materials and Methods

### Participants

Thirty-two native Spanish speakers were recruited for the current experiment and compensated €10 per hour for their participation. All participants had normal or corrected-to-normal vision and reported no history of medical, neurological, or psychiatric disorders. Two participants were excluded from the study due to technical problems during the EEG recordings. Data from 30 participants (17 females; age range 18–32 years, M = 23.77, SD = 4.38) were analysed. Informed consent was obtained from all participants in accordance with procedures approved by the Ethics Committee of the University of Barcelona.

### Stimuli

The experimental design included 312 images (350×350 pixels each): 104 images of famous faces (52 male and 52 female), 104 images of famous places, and 104 object images. Famous face and scene images were selected from a larger sample of the image database consisting of 284 and 184 pictures of each category, respectively. The selection was carried out by a separate sample of 10 Spanish university students (5 females; age range 21-39 years) who rated their familiarity with each image on a scale from 1 to 4 (1: Not recognised; 2: Familiar; 3: Recognised but don’t know the name; 4: Know the name). The final set of 104 face and place images were those that received the highest mean score from 10 external raters (mean score equal to or higher than 3.44 for males, 2.89 for females, and 2 for places). The 104 objects were selected from available object-picture databases and covered 6 categories (clothing, food, tools, transport, work, and leisure). Among the 312 images, 60 images (20 object images, 20 face images of famous people, and 20 images of famous places) were randomly selected for the localiser phase. For the main task, 36 images (12 object images, 12 face images, and 12 place images) were used for example trials and the rest 216 images (72 object images, 72 face images, and 72 place images) were used for the encoding trials; this separation was kept the same across participants.

### Experimental design

The experiment consisted of the localiser phase and the task phase. In the localiser phase, 60 images (20 faces, 20 scenes, and 20 objects) were presented in random order to participants. Each trial started with a 1000 ms fixation cross, followed by a 2500 ms image presentation. A text displayed on the screen then indicated the need for the participants to state the category of the just-presented image (**Figure 1**). Participants had a maximum of 10 seconds to respond. The next trial started immediately once a response was given or the maximum time limit was passed. There was a brief break between every 20 trials when participants could briefly rest and decide to continue whenever they felt ready.

**Figure 1.**
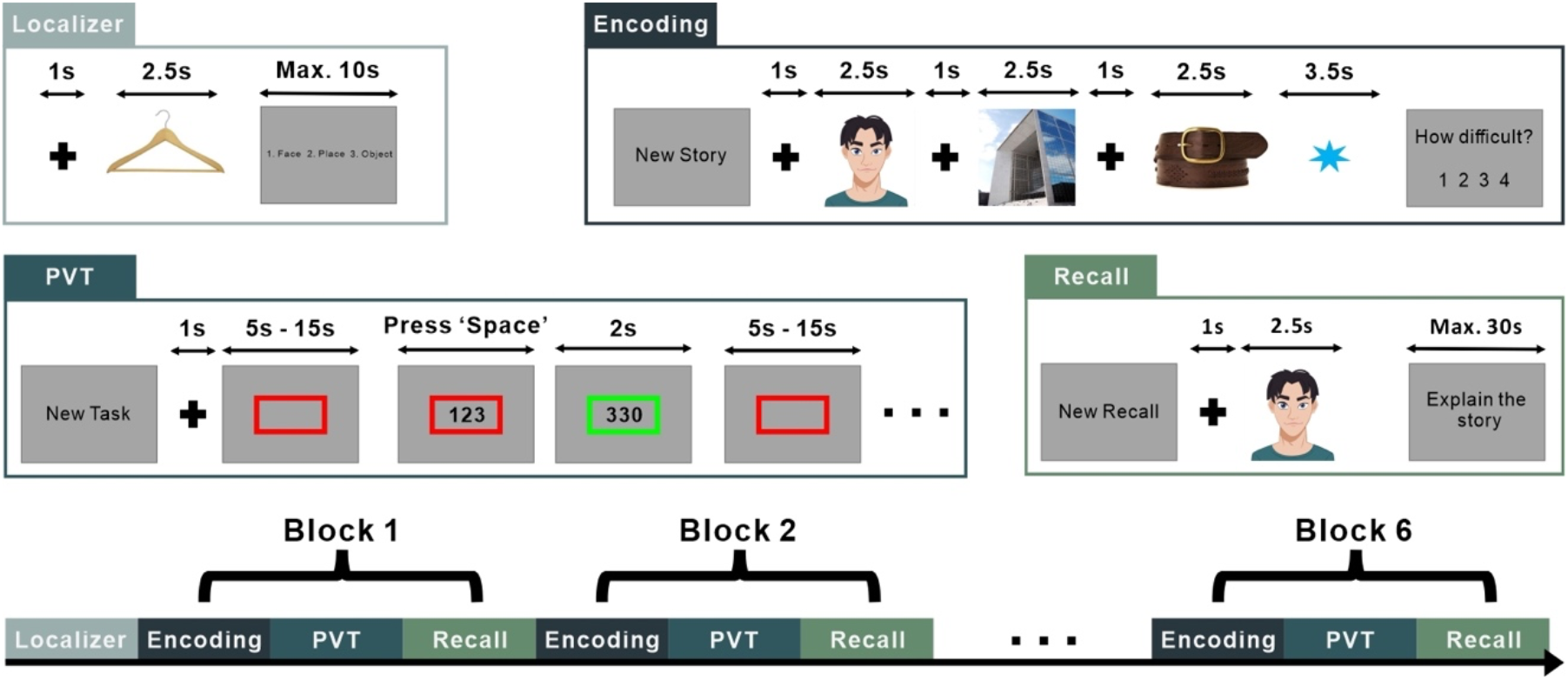
Experimental design. For the localisation task, 60 different images from 3 categories (face, place, and object) were presented. Participants were asked to indicate the image category. The encoding phase consisted of an image of an object, a famous face, and a famous place. Participants were asked to construct stories using the three elements for a later memory test. A blue asterisk appeared at the end of each triplet and participants rated their subjective sense of difficulty for story construction. For the Psychomotor vigilance task, participants were instructed to pay attention to the centre of the screen, waiting to react at the onset of a text timer by pressing the ‘Space’ button. In the recall phase, there were 12 recall trials, each of which used the first image of the previously presented triplets to cue the free recall of the other two images. One block was completed after the retrieval phase and the next block started following a brief break. The experiment consisted of 6 blocks in total. An avatar image is displayed due to bioRxiv policy on not displaying pictures of real people.

The task phase started after the localiser phase. The task phase included 6 blocks, each of them including an encoding task, a Psychomotor vigilance task (PVT), and a retrieval test. Each block was independent of the other, meaning that the images presented in one block were never shown in any other blocks. However, the task instructions and their order of alternation remained the same across all blocks. In the encoding task, participants were instructed to encode 12 series of three images, namely an object (O), a famous face (F), and a famous place (P). Participants were encouraged to construct stories using triplet elements in the form of a narrative (e.g., Iniesta went to Paris and purchased an expensive belt), and they were informed that the triplet information would be tested later. In total, 72 triplets were randomly generated for each participant from 216 images (72 object images, 72 face images, and 72 place images), and each image was used only once in the experiment. In each block, the presentation order of the image categories in a series was fixed (e.g., always ordered as Face-Place-Object in one block). There are in total 6 possible presentation orders, each of which was used in one of the 6 blocks with no repetition and randomly generated for each participant. At the beginning of each block, two example trials were presented, indicating the order of presentation of the image categories. Participants were instructed to use the two example trials to rehearse the upcoming series encoding in the block and told that the example trials would not be tested later. Each encoding trial began with the presentation of the text ‘New Story’ for 3000 ms, which marked the start of a new triplet series. Triplet images were then presented sequentially on a white screen for 2500 ms each after a 1000 ms black fixation cross. Immediately after the presentation of the last image in each triplet, a blue asterisk appeared on the screen, indicating a post-episode offset period of 3500 ms, during which participants were instructed to avoid rehearsing the just-encoded triplet series. Instructing participants not to rehearse the just-encoded episode served a dual purpose. Firstly, it helped minimize the active maintenance memory processes in the working memory. Secondly, it aimed to simulate naturalistic episodic memory formation circumstances where rehearsal is not a regular occurrence at the end of an episode. The asterisk remained visible on the screen during the offset period. Participants were then asked to provide a degree of subjective feeling of the difficulty of constructing a coherent episode with the just-presented triplet of images by a button press on a scale from 1 (‘very easy’) to 4 (‘very difficult’). The next trial began immediately after a response was given, or no response was given after a time limit of 10 seconds. A small break of ∼10 sec was provided after completing 6 trials.

A block of the PVT task followed the encoding phase. In each PVT block, participants were instructed to pay attention to the screen’s centre and press the space button as quickly as possible once the timer started counting. The task commenced with the text presentation ‘New Task’ for 3000 ms. Then an empty red square was displayed at the centre of the screen following a 1000 ms fixation cross. After a random interval of between 5 sec to 15 sec, the timer started counting in the middle of the square indicating the real passing time in milliseconds. The timer counted to a maximum of 3500 ms if no response was given. Once the participants pressed the button during the counting period, the timer stopped with the presentation of the final reaction time in the centre of the screen for 2000 ms. In cases where no response to the timer was given within the time limit, the presentation would be the final counting time of the timer (i.e., 3500 ms). The new PVT trial started immediately after the reaction time presentation. In total, 12 repetitions of response were required with no interruption in the middle. A block of a PVT task lasted around 3 minutes. The PVT task served as a distractor task adding a brief time interval between the encoding and the upcoming retrieval phase. It had the advantage of maintaining the attentive state of the participants and preventing them from actively rehearsing the just-encoded episodic sequences, and at the same time, it did not engage participants in any content encoding as the task is primarily reaction-based.

The PVT task was followed by a cued-recall task. During this task, participants were presented with the first image of all the encoded triplets in the current block in random order. They were required to verbally recall the story episode containing the other two images associated with the cue image. Each trial began with the text ‘New Recall’ for 3000 ms, followed by the cue image on the screen for 2500 ms and a 1000 ms fixation cross. The text ‘Explain the story’ was then displayed on the screen, which indicated to the participants they could start the verbal recall. The verbal recall had a maximum duration of 30 s, during which the text instruction remained visible on the screen all the time. Participants could skip to the next trial when finished with their recall or if they were unable to recall any associated image by pressing the space bar. A brief break of ∼20 s separated the start of the next block.

### EEG recording and pre-processing

During the experiment, EEG was recorded with a 64-channel system at a sampling rate of 512 Hz, using a eego™ amplifier and Ag/AgCl electrodes mounted in an electrocap (ANT neuro) located at 59 standard positions (FP1/2, AF3/4, Fz, F7/8, F5/6, F3/4, F1/2, FCz, FT7/8, FC5/6, FC3/4, FC1/2, Cz, T7/8, C5/C6, C3/4, C1/2, CPz, TP7/8, CP5/6, CP3/4, CP1/2, Pz, P7/8, P5/6, P3/4, P2/1, POz, PO7/8, PO5/6, PO3/4, Oz, O1/2) and the left and right mastoids. Horizontal and vertical eye movements were monitored with electrodes placed at the right temple and the infraorbital ridge of the right eye. Electrode impedances were kept below 10 kΩ. EEG was re-referenced offline to the linked mastoids. Bad channels were interpolated, and a band-pass filter (0.5 Hz -30 Hz) was implemented offline. Blinks and eye movement artifacts were removed with independent component analysis (ICA) before the analysis.

### Behavioural data analysis

During the retrieval phase of the experiment, participants were instructed to verbally recall the constructed story episode associated with the picture cue. Verbal recall of each trial was recorded through an audio recorder, and the audio files were later analysed. A successful recall of the image was considered as either correctly mentioning the name or describing it in precise detail. Memory for each triplet was quantified by the number of images correctly recalled, namely 0, 1, or 2 images successfully recalled following the cue.

### EEG data analysis

For each participant, we extracted epochs of EEG activity surrounding pictures presented in the localiser and the encoding tasks. These EEG trial epochs had a duration of 2500 ms (1280 data points given the 512 Hz EEG recording sampling rate), and they were baseline corrected to the pre-stimulus interval (-100 to 0 ms). For later analysis, we focused on the first 2000 ms of the trial epochs from stimulus onset. We also extracted EEG epochs of 3500 ms (1792 data points) from the offset period following the encoding of each triplet series. EEG signal to the offset period was baseline corrected to the -100 to 0 ms averaged EEG activity. EEG trial epochs that exceeded ± 100 µV were discarded for further analysis. EEG trials were then Gaussian smoothed by averaging data via a moving window of 100 ms (excluding the baseline period) and then downsampled by a factor of 5.

### Representational Similarity Analysis (RSA)

RSA was performed timepoint-to-timepoint and included spatial features (i.e., scalp voltages from all 59 electrodes) (Silva et al., 2019; Sols et al., 2017; Wu et al., 2022). The similarity analysis was calculated using Pearson correlation coefficients, which are insensitive to the absolute amplitude and variance of the EEG response.

We conducted a trial-based RSA between the EEG signal elicited by each encoding item (1st, 2nd, and 3rd, regardless of the image category) and the EEG signal elicited at the offset period following the encoding of the triplet series. After smoothing and down-sampling, EEG epoch data elicited by each picture in the triplet included 205 sample points (given the 512 Hz EEG recording sampling rate) covering the 2000 ms of picture presentation and EEG data from post-triplet offset contained 359 time points, equivalent to 3500 ms. Point-to-point correlation values were then calculated, resulting in a 2D similarity matrix with the size of 205×359, where the x-axis represented the episodic offset time points and the y-axis represented the picture encoding time points. The output 2D matrix depicted the overall degree of similarity between EEG patterns elicited by each encoding image and the subsequent post-episodic offset interval.

We applied the same approach to explore the neural similarity between encoding items (1^st^, 2^nd^, and 3^rd^) in the sequence. To do so, we separately conducted RSA between the 1^st^ item and the 2^nd^ item, also between the 2^nd^ item and the 3^rd^ item. Point-to-point correlation values were then calculated on smoothed and down-sampled EEG epoch data, resulting in a 2D similarity matrix with the size of 205×205, where the y-axis represented the encoding time points of the items appeared earlier in the sequence (i.e., 1^st^ or 2^nd^ item) and the x-axis represented encoding time points of the items appeared later in the sequence (i.e., 2^nd^ or 3^rd^ item).

To account for RSA differences between conditions, we employed a nonparametric statistical method (Maris & Oostenveld, 2007), which identifies clusters of significant points on the resulting 2D similarity matrix and corrects for multiple comparisons based on cluster-level randomisation testing. Statistics were computed on values between conditions for each time point, and adjacent points in the 2D matrix that passed the significance threshold (*p* < 0.05, two-tailed) were selected and grouped together as a cluster. The cluster-level statistics took the sum of the statistics of all time points within each identified cluster. This procedure was then repeated 1000 times with randomly shuffled labels across conditions. Cluster-level statistics with the highest absolute value for each permutation were registered to construct a distribution under the null hypothesis. The nonparametric statistical test was calculated by the proportion of permuted test statistics that exceeded the true observed cluster-level statistics.

We also examined whether possible RSA effects (i.e., at cluster level) that could be seen when comparing successful and unsuccessful conditions in the previous analysis were trial-specific or whether they reflect task-specific patterns of correlation between online and offline encoding time periods. In other words, we aimed to assess whether RSA in the same trial between image encoding and offset period from the same trial was higher than RSA between image encoding and offset periods from different trials. To assess these issues statistically, we ran the RSA individually and separately for successful and unsuccessful memory conditions by randomly shuffling the paring of EEG data from a given triplet and an offset period. This procedure was repeated 200 times, each with a randomly generated shuffling order. The results were then averaged across permutated trials for the identified cluster and compared to the real cluster value using a repeated-measure ANOVA.

### Linear Discriminant Analysis (LDA)

To identify the multivariate pattern of brain activity for image processing of different categories, a Linear Discriminant Analysis (LDA) was trained and tested on the EEG sensor patterns of localiser trials (pre-processed signal amplitude from 59 channels). The classifier was trained independently per participant and at each time point during localiser image presentation, then tested with a leave-one-out cross-validation procedure. Given that three categories were included in the current experiment (face, place, and object), at the training stage, the classifier was trained repetitively three times, including each possible pair out of the three classes. For each of the two classes, the classifier found the decision boundary that best separated the pattern activity. We then asked the classifier to estimate the unlabelled pattern of brain activity for each of the three decision boundaries (one for each pair of classes). The output of the classifier for each two trained classes at a given time point was the distance value to the decision boundary, which represents how probable the pattern of brain activity belonged to one of the two included classes, with the sign indicating the class and the magnitude reflecting the confidence of the classifier. The distance value for each pair of classes was then sigmoid transformed to get the probability of either class that unlabelled pattern activity belonged to (e.g., a distance value of 0 will return 50% for either class). After normalising and averaging values across the three possible pairings, the class with the highest probability was marked as the final label for the testing data. To access the general separability between the three classes in a compound measure, we defined a separability index (*D* value) as the sum of the absolute of the three distance values to each of the decision boundaries (i.e., used to threshold the classification decision for each pair of classes), with the assumption being that the greater the separability index, the higher the probability that the given activity pattern belonged to a specific class rather than assimilating to all three classes with equal distinctiveness (i.e., closer to zero).

This training-test procedure was repeated until every single localiser trial had been classified. The predicted labels for all trials at every given time point were then compared to the true classes to assess the accuracy of the classifier across all localiser times.

To evaluate how face, object, and scene category representations accounted for EEG patterns elicited during picture encoding and at the offset period, we first identified the time point where the cross-validation of the classifier reached the peak accuracy. Using patterns of activity surrounding 10 time points around the peak (-50 ms to 50 ms with the peak time point in the middle), we then trained the classifier per participant with all localiser trials and predicted all sample points separately for encoding and offset trials. The results were then averaged across localiser time points, resulting in a 1D separability index (*D* value) line for each trial where each sample point represented the encoding/offset time points.

Note that for all the LDA analyses in this study, the training set and test set for each participant were z-transformed for each channel and each timepoint across all the trials before the application of LDA.

### Linear-mixed effect model

To further explore how the separability of pattern activity between picture categories changed along the encoding sequence and whether it is predictive for behavioural memory on a trial basis, we implemented a Linear Mixed Effect Model (LMM) on the resulting general distance value of each encoding image classified by patterns trained on trials from the localiser task. We further smoothed the resulting 1D distance value for each predicting encoding trial by averaging over a moving window of 200 ms, then introduced in our LMM the *D* value at each time point as the dependent variable, then both the image order in triplet series (1^st^, 2^nd^, and 3^rd^) and recall memory (successfully recalled 2 images or not following the cue), as well as the interaction of the two as fixed effect variables. Subject was introduced in the model as the grouping variable, with random intercept and a fixed slope for each fixed effect variable. The statistical significance was then evaluated using Bonferroni correction for each fixed effect variable at each timepoint thresholded with an adjusted alpha level of α = 2.44×10^−4^ (0.05/205).

## Results

### Localisation task

For the localisation task, 26 out of 30 participants reached 100% accuracy in identifying the image category, and the mean accuracy across the 30 participants was 99.72% (SD = 0.77%).

### Recall of picture triplets

Participants were able to recall on average 1.12 pictures (SD = 0.37) following the cue, with the mean percentage of trials recalling 0, 1, and 2 items being respectively 35.61% (SD = 16.41%), 16.30% (SD = 6.52%) and 48.09% (SD = 20.68%) (**Figure 2a**). Separated by the category of item, we found that face as a to-be-recalled item (i.e., not the first picture in the triplet series) was better remembered (Mean = 62.15%, SD = 17.70%) compared to place (Mean = 51.84%, SD = 19.98%) and object (Mean = 54.70%, SD = 19.25%)(repeated measures ANOVA: *F*_(2,58)_ = 24.379, *p* < 0.001). Relatedly, there was a significant difference in recall performance depending on the category order of the triplet series. More concretely, we found that encoding blocks that included triplets with face as the first picture (i.e., Face-Place-Object or Face-Object-Place) were less accurately recalled (Face-Place-Object: mean = 0.914, SD = 0.098; Face-Object-Place: mean = 0.905, SD = 0.082) than triplets from blocks where place (Pace-Face-Object: mean = 1.253, SD = 0.081; Pace-Object-Face: mean = 1.224, SD = 0.081) or object (Object-Face-Place: mean = 1.213, SD = 0.081; Object-Place-Face: mean = 1.230, SD = 0.084) was presented first (repeated measures ANOVA: *F*_(5,140)_ = 9.798, *p* < 0.001). We also found that the average number of images recalled upon the picture cue did not vary between blocks, indicating that encoding and retrieval accuracy did not vary throughout the task (repeated-measures ANOVA with block number as the main factor: *F*_(5,140)_ = 0.415, *p* = 0.838).

**Figure 2.**
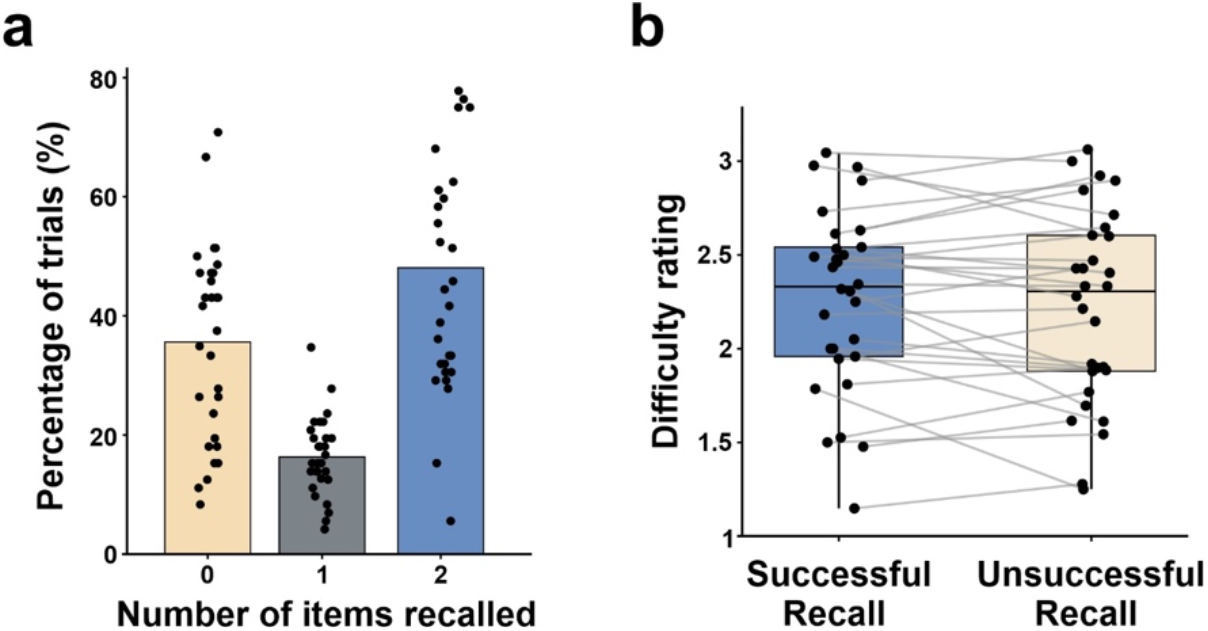
Behavioural results. **(a)** Percentage of trials as a function of numbers of images correctly retrieved during free recall. **(b)** Subjective rating of the difficulty of triplet encoding separated by whether or not the triplet was later successfully recalled (with both images associated with the cue being correctly recalled). Each dot on both plots represents the value for an individual in the corresponding condition. Each grey line on the boxplot connects the value of an individual in two conditions.

For RSA analysis, we adopted a median-split approach to separate the trials based on whether the entire triplet images were correctly retrieved. Triplets with 2 images recalled after the cue were labelled as successful recall, and triplets with either 1 image or no image recalled were labelled as unsuccessful recall. The average percentage of trials was respectively 48.09% (SD = 20.68%) for successful recall condition and 51.91% (SD = 20.68%) for unsuccessful recall condition (Wilcoxon signed-rank test: *z* = -0.43, *p* = 0.67).

### Participants’ ratings of encoding difficulty

On average, triplets were rated as 2.24 (SD = 0.47) (on a scale that ranged from 1: no difficulty to 4: very difficult), and the mean percentage of triplets rated as 1, 2, 3, and 4 were respectively 27.91% (SD = 21.73%), 33.79% (SD = 13.80%), 24.65% (SD = 13.13%) and 13.65% (SD = 12.03%). Based on the median-split criteria, difficulty ratings for trials with successful recall (mean = 2.26 and SD = 0.48), and for trials with unsuccessful recall (mean = 2.22 and SD = 0.51) did not differ statistically between each other (paired Student *t*-test: *t*_(29)_ = 1.01, *p* = 0.32, two-tailed) (**Figure 2b**).

### RSA between item sequence and episodic offset at encoding

We first examined the existence of encoding-offset neural similarity differences between trials that were successfully or unsuccessfully recalled. This analysis revealed that EEG patterns elicited during the encoding of picture triplets that were later recalled showed, compared to unsuccessfully recalled trials, a higher degree of neural similarity during the episodic offset period (**Figures 3a and b**). This result was corroborated statistically with the cluster-based permutation test, which showed one cluster of increased neural similarity starting at ∼200 ms at offset period (*p* < 0.001 (corrected), mean *t*-value = 2.45, peak *t*-value = 4.92) (**Figure 3b**). We next evaluated encoding-offset neural similarity corresponding to each element in the sequence. We extracted the mean similarity values within the identified cluster for each item–offset pair and computed a repeated-measures ANOVA with two factors: Trial Condition (successful vs. unsuccessful recall) and Item Order (1^st^, 2^nd^, and 3^rd^ in the sequence). The results of this analysis showed a significant main effect for Trial Condition, *F*_(1, 28)_ = 19.171, *p* < 0.001, and a marginally significant main effect for Item Order (*F*_(2, 56)_ = 3.047, *p* = 0.055), and no interaction between the two main effects (*F*_(2, 56)_ = 0.368, *p* = 0.694). Pairwise comparisons showed marginally significantly higher mean cluster similarity values for 3^rd^ item in the sequence (mean = 0.028, SD = 0.011) as compared to the 1^st^ (mean = 0.012, SD = 0.008, *p* = 0.05) and 2^nd^ item (mean = 0.011, SD = 0.009, *p* = 0.053) (**Figure 3c**).

**Figure 3.**
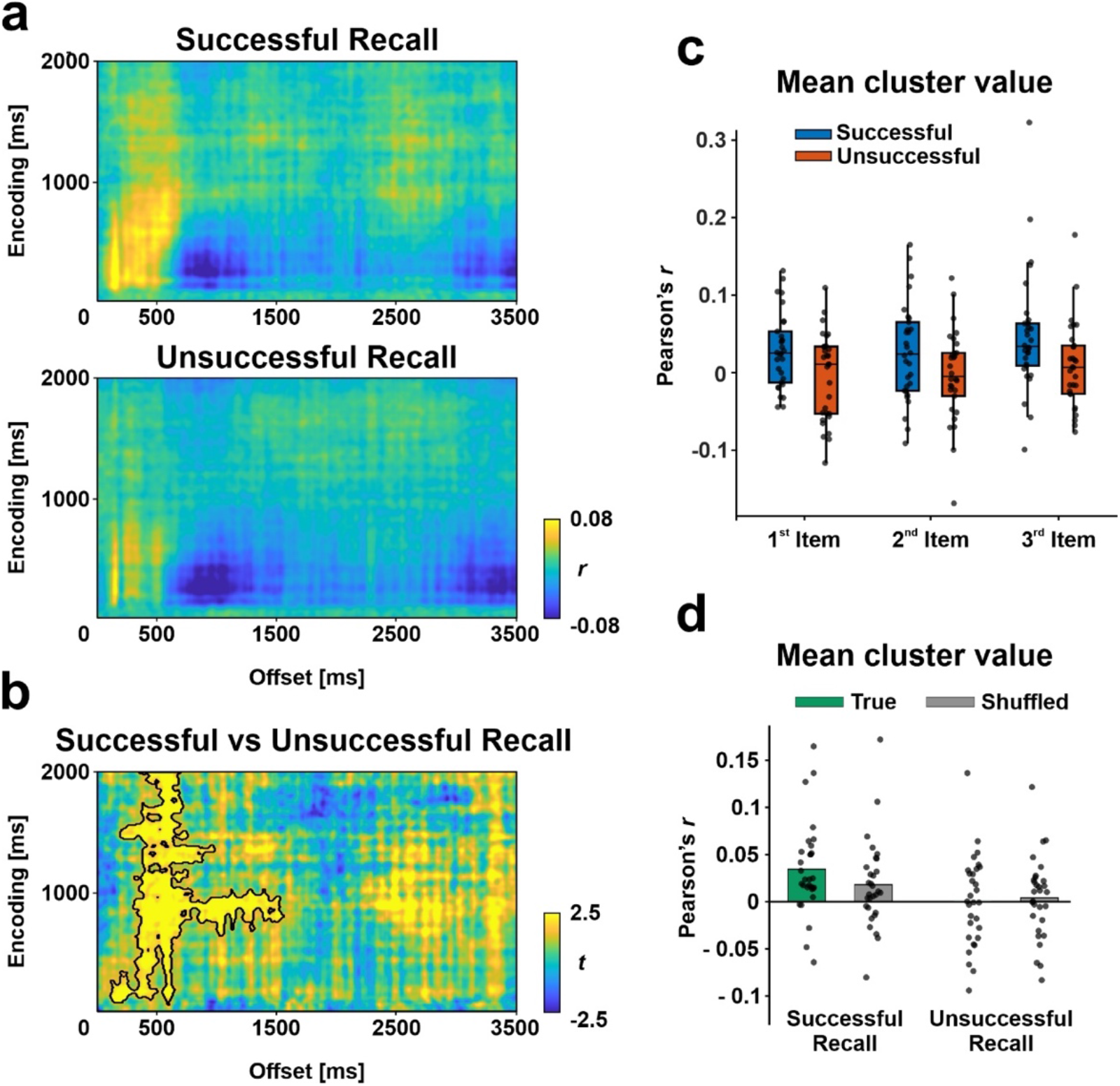
RSA for image at encoding and post-encoding. **(a)** Time-resolved degree of neural similarity between image encoding and post-triplet offset for trials with successful subsequent recall (upper) and unsuccessful recall (lower). **(b)** Difference between similarity values for the two conditions. Statistically significant (*p* < 0.05, cluster-based permutation test) higher similarity value was found for trials with successful recall centred in one area (indicated by black contour lines). (**c**) Averaged RSA values within the cluster separated by successful and unsuccessful subsequent recall for each encoding item. The central mark is the median across participants and the edges of the box are the 25th and 75th percentiles. Each black dot represents values for an individual participant. **(d)** Averaged true RSA vs. shuffled RSA values within the cluster separated for trials with successful and unsuccessful subsequent recall. Each dot on the plots represents the value for an individual in the corresponding condition.

We next tested whether the RSA effects seen when comparing successful and unsuccessful conditions were trial-specific. To address this issue, we ran RSA by shuffling the encoding-offset pairing 200 times and obtained for each individual a similarity value that represented task-rather than trial-specific RSA effects. To evaluate whether real and shuffled RSA effects differed from each other statistically, we conducted a repeated-measure ANOVA with two main factors, Trial Condition (successful vs. unsuccessful recall) and RSA condition (true vs. permutated value). We found a significant main effect for Trial Condition (*F*_(1,29)_ = 16.47, *p* < 0.001) and RSA condition (*F*_(1,29)_ = 4.75, *p* = 0.037), as well as the interaction of the two main factors (*F*_(1,29)_ = 12.41, *p* = 0.001) (**Figure 3d**). These results indicated that the RSA showed a trial-specific property only for successfully remembered trials.

### Classification accuracy and separability of picture category

We adopted the LDA approach to classify and predict the image category being processed based on the elicited EEG pattern in the localiser task. The classifier was trained independently per participant and at each time point during picture encoding, then tested with a leave-one-out cross-validation procedure. Two output values were extracted for each time of training/testing, namely the category of tested data predicted by the model with the highest probability among three alternatives (i.e., accuracy) and a general distance value (*D* value) of tested data to the classification plane among categories (i.e., separability) (**Figure 4a**).

**Figure 4.**
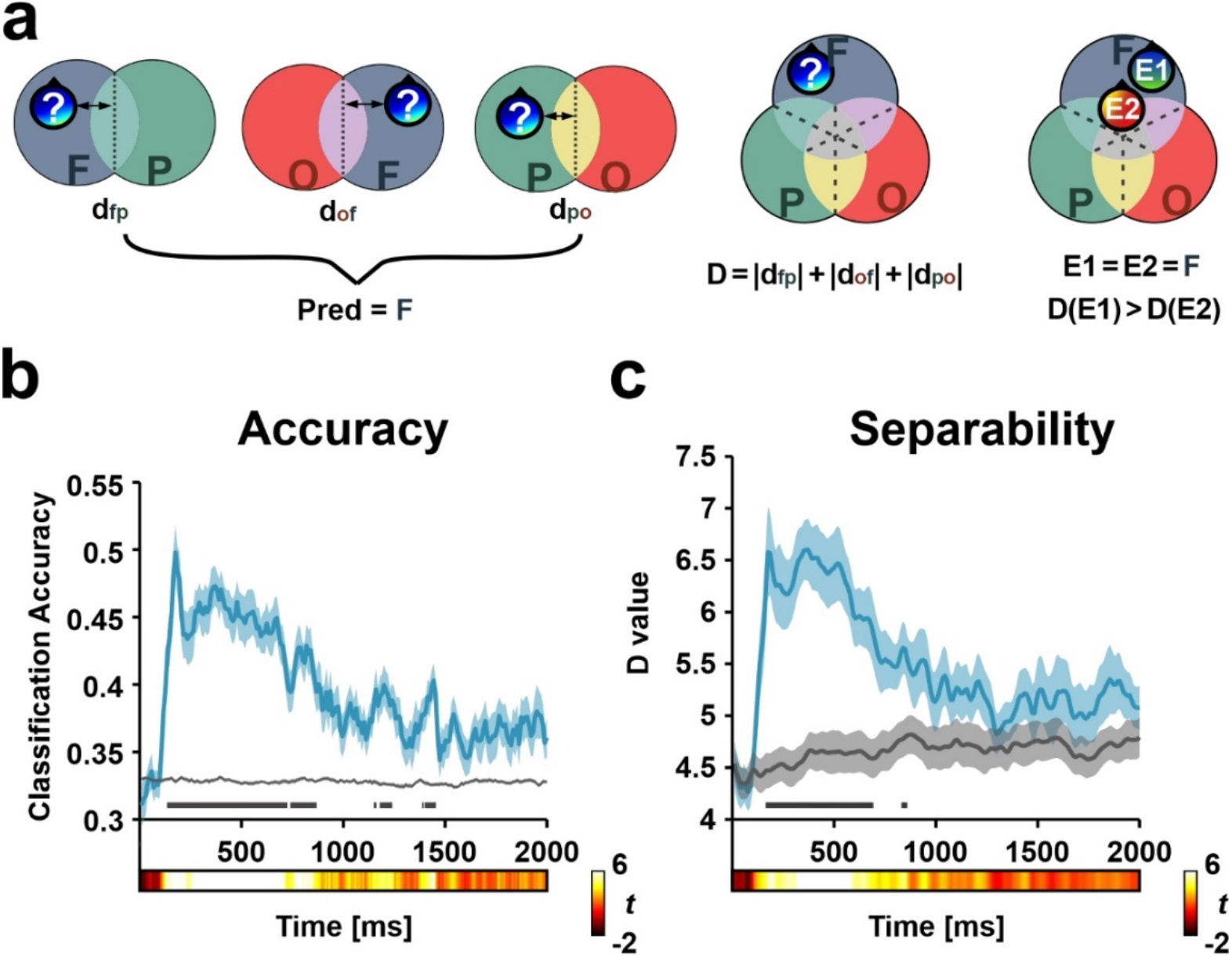
Two cross-validated LDA classifier output measures using localisation trials. **(a)** Abstract illustration of the classifier output calculation. The distance value for each pair of classes was sigmoid transformed to get either class’s probability. The class with the highest probability after normalising and averaging values across three pairs was marked as the final label for the testing data (left). The general distance value (*D* value) was defined as the sum of the absolute of the three distance values to each of the decision boundaries (middle). Pattern examples 1 and 2 can be both classified accurately as ‘Face’ images. However, Pattern example 2 also showed a more similar pattern to the ‘Place’ and ‘Object’ categories, which a smaller *D* value can indicate. **(b)** Classifier accuracy estimated using the leave-one-out method. Image categories can be reliably decoded compared to surrogate trials starting around ∼130 ms after image onset, with peak value reaching ∼180 ms. **(c)** Pattern separability among image categories quantified by the general distance value, which evolved similarly across time compared to the accuracy measure. *D* value reached the peak at ∼180 ms and ∼380 ms. In plots (***b***) and (***c***), the shaded area indicated SEM across participants, and statistical significance compared to surrogate trials was Bonferroni-corrected and marked in dark grey line.

The results of this analysis showed that picture category could be reliably predicted rapidly at picture onset (i.e., ∼130 ms), showing a peak classification accuracy at ∼180 ms (*t*_29_ = 8.41, *p*_corr_ < 0.001) (**Figure 4b**).

As expected, the pattern separability analysis showed similar temporal dynamics to the accuracy ones. More specifically, pattern separability became significant as the distance value increased compared to surrogate trials, with the difference emerging from ∼170ms. *D* value reached the local maximum at respectively ∼180 ms (D = 6.57, *t*_29_ = 5.032, *p*_corr_ = 0.005) and at ∼380 ms (D = 6.60, *t*_29_ = 7.25, *p*_corr_ < 0.001) (**Figure 4c**).

### Gradual integration of picture category information during sequence encoding

We examined whether the sequential encoding of pictures from different categories in the encoding task would involve a gradual integration of the just-encoded images from the sequence and whether this process predicted memory recall. To address this issue, we extracted the -50 to 50 ms EEG pattern surrounding the peak (i.e., at 180 ms from picture onset; **Figure 4b**) LDA accuracy during the encoding of images in the localiser task. We then used these EEG patterns as the training data in a new LDA and tested on EEG patterns elicited at each time point from each picture from the sequence on the encoding task.

We then averaged across all training time points at trial level and included the resulting distance value at each time point of encoding into LMM as the dependent variable. For each trial, the memory condition (successful vs unsuccessful recall), the encoding order of the image in the triplets (i.e., 1^st^, 2^nd,^ and 3^rd^), and the interaction of the two were included in the model as fixed-effect variables. Subject was introduced into the model as the grouping variable, with random intercept and a fixed slope for each fixed-effect variable.

This analysis showed that the *D* value correlated negatively with the order of picture in the sequence (**Figure 5a**) and that such effect emerged during ∼ 460ms - 520ms and ∼590ms - 920ms after picture onset. However, we found that *D* value did not correlate with later trial memory at the test nor the interaction of picture order and memory. This suggested that the picture category integrative process takes place during sequence encoding and that this had no impact on the later ability of the participants to retrieve the sequence episode. Different from the *D* value, the classification accuracy showed a similar pattern as a function of the order of pictures in the sequence (**Figure 5b**). To control for the possibility that the observed effect was not merely due to a decrease in the specific category classification accuracy as a function of the order of the picture in the sequence, we extracted the mean accuracy across image order within the time window where the significant decrease in pattern separability was identified. A repeated-measure ANOVA showed significantly above-chance accuracy value (*F*_(1,29)_ = 32.4, *p* < 0.001) with no main effect for image order (*F*_(2,58)_ = 0.004, *p* = 0.99) (**Figure 5c**). Jointly these results indicate that while the classifier continued to predict the image category accurately, the decrease in the *D* value as the triplet series unfolded reflects a gradually reduced pattern separability among categorical representations during image encoding, suggesting an integration process associating the upcoming stimulus representation to the previous ones.

**Figure 5.**
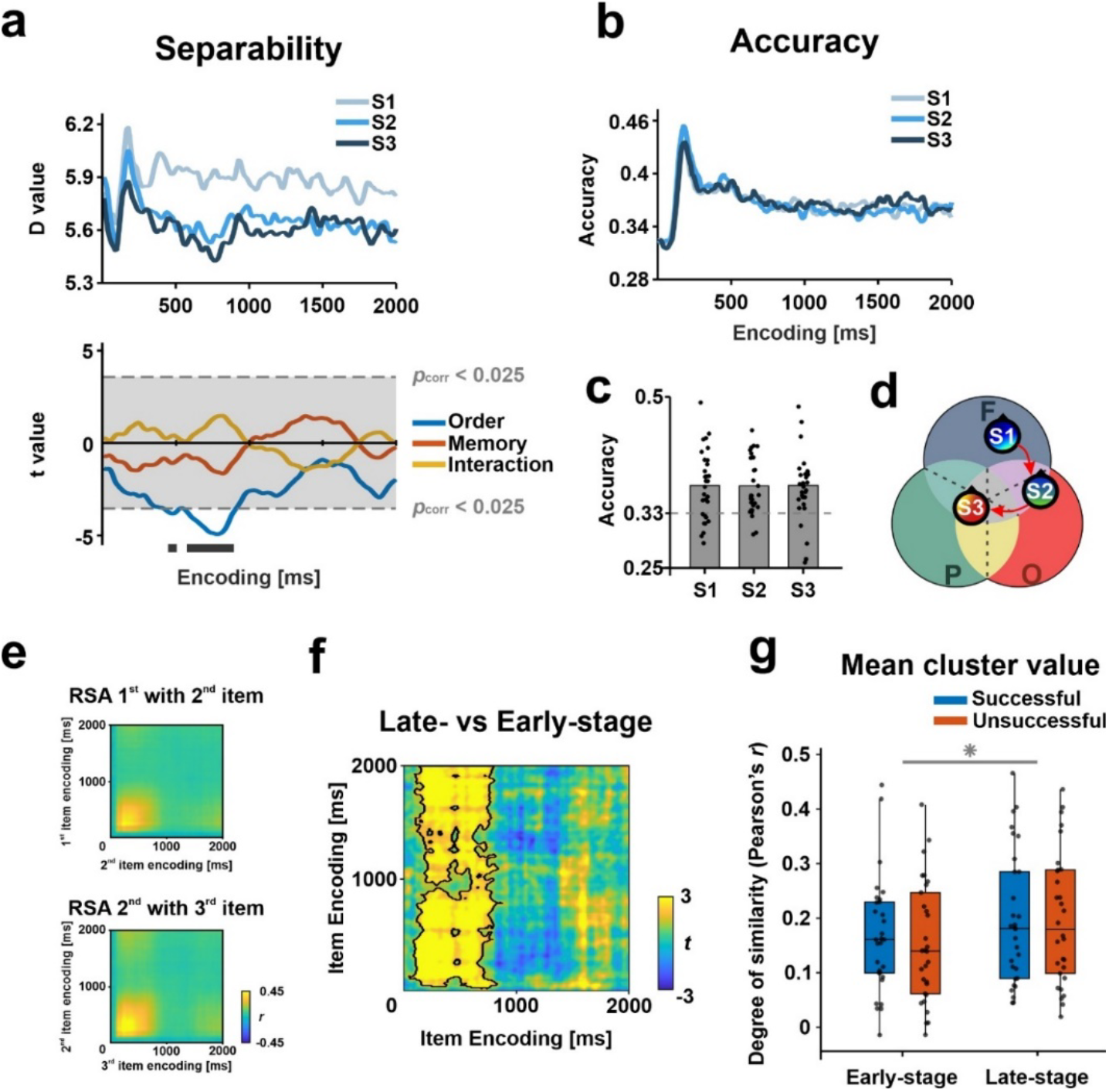
Representational dynamics and transformation during sequence encoding. **(a)** Across participants averaged *D* value (i.e., degree of category separability) during encoding as a function of the order of image in the sequence (upper) with statistical significance Bonferroni-corrected for the main effect of image order, subsequent memory, and their interaction (lower). Shaded grey area and light grey dashed line marked the significance threshold boundaries (two-tailed) adjusted by Bonferroni correction. Time window where the main effect passed the threshold was marked in dark grey line below. **(b)** Across participants averaged LDA accuracy during encoding as a function of the order of pictures in the sequence. **(c)** LDA accuracy averaged across the time window where a significant effect for image order was found. Category of image was classified equally accurate across image order (*p* = 0.99) yet significantly above chance (grey dashed line) (*p* < 0.001). Each black dot represents values for an individual participant. **(d)** Abstract illustration of the speculated integration process. While the classifier continued to predict the image category accurately, there was a trend for an ‘integrated’ pattern indicated by a gradually decreased pattern separability. **(e)** Time-resolved degree of neural similarity between encoding items for 1^st^ with 2^nd^ item (upper) and 2^nd^ with 3^rd^ item(lower). **(f)** Difference between similarity values for late-stage items pair (2^nd^ with 3^rd^) vs early-stage item pair (1^st^ with 2^nd^). Statistically significant (*p* < 0.05, cluster-based permutation test) higher similarity value was found for late-stage items pair centred in one area (indicated by black contour lines.). **(g)** Mean similarity values within the identified cluster between encoding items. Each black dot represents values for an individual participant. Late-stage items pair in the sequence (2^nd^ with 3^rd^) showed significantly greater similarity as compared to early-stage items pair (1^st^ with 2^nd^). The central mark is the median across participants and the edges of the box are the 25th and 75th percentiles.

Having shown that categorical information is gradually integrated during encoding, we aimed to investigate the potential for unique and individual representations to evolve over time. To achieve this goal, we conducted RSA between the encoding items within the sequence (as described in the Methods section) (**Figure 5e**). Our findings revealed that neural similarity between the encoding items was greater for the item pair that emerged later in the sequence (i.e., 2^nd^ with 3^rd^) in comparison to the item pair that emerged earlier in the sequence (i.e., 1st with 2nd) (*p* < 0.001, corrected by cluster-based permutation, mean *t*-value = 3.067, peak *t*-value = 8.141) (**Figure 5f**). These results supported the notion of a gradual formation of an integrated representation as the sequence progressed. To further assess whether the increase in between-item similarity as the sequence unfolded was linked to memory performance at test, we calculated the mean similarity values within the identified cluster for each of the two between-item pairs, separately for trials that were successfully and unsuccessfully recalled at test. We conducted a repeated-measures ANOVA on the mean similarity values within the cluster across participants, with two factors: Encoding Stage (early vs. late) and Trial Condition (successful vs. unsuccessful recall). Our analysis revealed a significant effect for Encoding Stage (*F*_(1,29)_ = 19.38, *p* < 0.001), but no effect for Trial Condition (*F*_(1,29)_ = 0.53, *p* = 0.472), nor a significant interaction between the two factors (*F*_(1,29)_ = 0.82, *p* = 0.372) (**Figure 5g**). These results suggest a gradual integration of idiosyncratic representations at the item level during encoding, similar to the integration process indicated by LDA for categorical information. Furthermore, this integration process does not directly influence how effectively these items will be retrieved at test.

### Picture sequence integration and memory at episodic offset period

We next examined whether an integrated form of the just encoded sequence could predict memory for the episode right after its online encoding, that is, once the encoding ended, at the offset period. If this was the case, we should observe that *D* value was reduced at the offset period for successful compared to unsuccessful recalled picture sequences.

To address this issue, we again extracted the EEG pattern elicited during ∼130 ms - 230 ms (-50 to 50 ms surrounding the peak classifier accuracy at 180ms from picture onset) by picture presentation in the localiser task. We applied it to each time point of the post-episodic offset period. The resulting *D* value from the classifier was then averaged across all training timepoints. We further smoothed the resulting 1D distance value trial by averaging over a moving window of 200 ms, then separated *D* values for successful and unsuccessful memory trial conditions and averaged them for each participant. Statistical comparisons between conditions were assessed and then reassessed with a cluster-based permutation approach.

The analysis revealed that the *D* value remained stable during the offset period, as depicted in **Figure 6**. Cluster-based statistics did not pinpoint a specific time frame in which successfully recalled trials differed significantly from unsuccessfully recalled ones. This suggests that the categorical representation formed during encoding is not enhanced at offset, and that the level of categorical information integration is not linked to memory performance.

**Figure 6.**
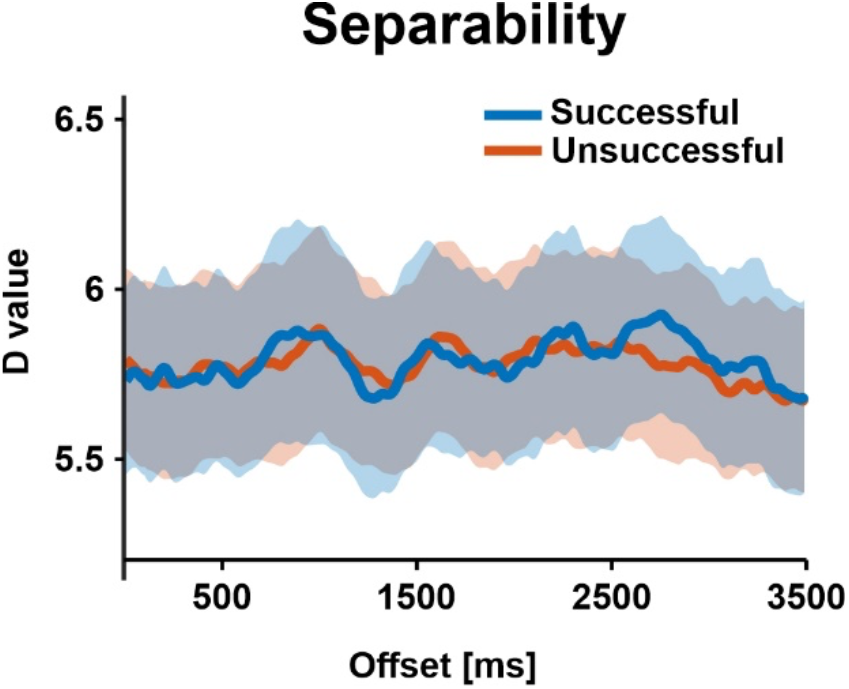
Pattern separability during post-triplet offset period predicted by LDA classifier trained on localisation trials. There is a consistent level of separability between categories throughout the offset period. Cluster-based statistics do not show a time window in which a statistically significant difference between successful and unsuccessful trials is found. Shaded area indicated SEM across participants.

## Discussion

The current study examined the dynamics of the representational format that contributed to successful episodic memory formation. The combination of RSA and multivariate decoding approaches on EEG data allowed us to systematically compare whether “category-level” or “item-level” representations supported memory formation during the online encoding of a picture triplet sequence and offline, in the period that immediately followed encoding. Our findings revealed a gradual integration of category-level representation during the online encoding of the picture sequence and a rapid item-based neural reactivation of the encoded sequence at the episodic offset. However, we found that only memory reinstatement at episodic offset was associated with successful memory retrieval from long term memory, thereby indicating that post-encoding memory reinstatement is akin to the rapid formation of unique memory for episodes that unfold over time.

Research in the field of visual object recognition has shown that perception involves dynamic transformations of information from low-level visual inputs to higher-level visual properties and ultimately complex semantic representations (Kravitz et al., 2013). This research has revealed that incoming information flows through lower-to higher-order brain regions during encoding, with more abstract information represented in the higher-order areas. Meanwhile, the higher-order regions interact with the lower-level regions to reshape the representations according to task requirements or assimilate them into pre-existing knowledge representations (Xue, 2022). Interestingly, it is widely accepted the encoding of the different representational formats occurs rapidly but temporally organized in the brain, so that visual object recognition starts with low-level perceptual followed by high-level abstract processing (Carlson et al., 2013; Cichy et al., 2014; Lehky & Tanaka, 2016; Serre et al., 2007). Our findings are consistent with previous research that relied on the implementation of pattern classification algorithms on EEG or MEG data in showing that category level representations can be detected at early time windows from image presentation onset (Cichy et al., 2014; Jafarpour et al., 2014; Wimmer et al., 2020). Our results contributed to this literature by showing that category-level representation associated with picture items in a sequence can be accurately detected with consistent time intervals and precision throughout the sequence.

Our findings contribute to our understanding of how category level representations associated with items from a sequence are integrated as the sequence unfolds. Concretely, we found a gradual process of integrative encoding of category-level representation. Specifically, we observed a gradual process of integrative encoding of category-level representations. The category representation of the second picture was found to be more similar to the category representation of the first item, while the category representation of the third picture integrated category-level representations from both the preceding second and first pictures in the sequence. This observation was quantifiable in our study because instead of registering only the output from the classifier, generally defined as the predicted class with maximal likelihood among possible alternatives, we used it to develop an index that quantified the classifier’s ability to distinguish among all the possible classes at a given time point, the degree of separability or the *D* value. In other words, the *D* value expresses the degree to which a tested neural pattern assimilated or deviated from all the possible trained categories. It is important to note that the gradual decrease in pattern separability as a function of picture order in the sequence was not accompanied by changes in pattern classifier accuracies. Thus, the observed gradual decrease in pattern separability cannot simply be attributed to a weak classification performance. It should be noted that the different interpretations drawn from the two measures of the classifier, namely accuracy and D value, are based on their distinct metrics. The accuracy measure refers to the prediction output of the classifiers as a categorical index, which depends on the relative comparison between the introduced classes, with the class having the largest distance to classification hyperplane being the winner. However, from a more general aspect, it does not provide any information about how close the to-be-predicted data to other classes’ neural patterns are. By summing up the absolute values of the distance to all the introduced class neural patterns, the *D* value provides a quantitative measure of how distinct in general the introduced data pattern is to all the classes, sensitive to the global changes in brain activity with respect to all the categories during encoding, a speculation the current study was particularly interested in.

One plausible neurobiologically-based explanation of the gradual decrease in pattern separability as a function of picture order in the sequence may be attributed to an attenuated neural activity by prior expectation, given that participants could anticipate the category of the upcoming image since the order of presentation was fixed within each block. Though prior studies revealed that anticipation might reduce response in neurons tuned for expected stimulus (Kok et al., 2013; Kumar et al., 2017), multivariate approaches have instead shown a ‘sharpening’ effect for perceptual representations in cortical regions due to a more selective population response (de Lange et al., 2018), resulting in a more accurate pattern classification (Kok et al., 2013). Our findings that the decrease in picture category pattern separability is taking place at around 500 ms from picture onset, however, may not be explained by ‘sharpening’ effects because they are thought to occur earlier in the temporal course of processing (i.e., < 400 ms from stimuli onset). Instead, we argue that the gradual reduction in pattern separability (i.e., decrease in the *D* value) but preserving the category-based discriminative accuracy (i.e., similar pattern classification accuracy) following the sequential presentation of images reflected a continuously additive category-specific processing, which promoted the encoding of multiple categorical information in parallel, supported by various overlapping cortical regions. In fact, different yet overlapping cortical regions (e.g., various regions on the lateral surface of occipitotemporal cortex) are selectively sensitive to stimuli from different categories when presented in isolation, including face, objects and scenes (Silson et al., 2016). In naturalistic scenarios, the processing of multiple categories of information embedded in the encoding experience takes place simultaneously, and the neural signature of such processes can be decoded in different cortical regions (Cooper & Ritchey, 2020). In the context of our study, the ongoing need to associate each appearing picture with the previously encoded pictures from the sequence may have promoted integrative processes online during the encoding of the picture.

We found no significant relationship between the degree of integration of item- or category-based representations during encoding and their later retrieval. However, we did find that the extent to which item-based information was reactivated at the episodic offset, just after the encoding of the sequence was completed, was a reliable predictor of a participant’s ability to recall the episodic trace from long-term memory. These results are in line with fMRI literature using different input types such as picture sequences (DuBrow & Davachi, 2014), short video clips (Ben-Yakov et al., 2013; Ben-Yakov & Dudai, 2011; Zacks et al., 2001), and movies (Baldassano et al., 2017; Ben-Yakov & Henson, 2018) that highlighted the sensitivity of the hippocampal-neocortical system to detect episodic offsets and its link to memory formation. In addition, the present findings add additional support to scalp EEG evidence of memory reactivation of just-encoded events upon the completion of an event supports successful memory formation (Silva et al., 2019; Sols et al., 2017; Wu et al., 2022). The current data however extends them by uncovering the representational format of this offset-locked neural activity. More specifically, our data suggests, consistent with studies on rodents (Foster & Wilson, 2006), that memory formation of single-trial sequential episodes is facilitated by the reactivation of high-fidelity representations of the encoded episodic sequence in memory. These results contrast with human studies, such as Liu et al. (2021), which found that memory reactivations during delay periods involving short-term memory maintenance transform the original representation into a more semantic representational format. Notably, while participants in our task were instructed to rest and refrain from rehearsal, participants in Liu et al. (2021) were actively maintaining the encoded information during the delay period. An interesting hypothesis to explore in future research would be that neural reactivation immediately after encoding can flexibly adjust the representational format of the replayed information in accordance with task goals. This would be consistent with recent findings in humans, which showed that the representational structure of replayed memories during a task shifted depending on whether the task required planning or preservation of the just-encoded information (Wimmer et al., 2023).

In conclusion, we found a gradual integration process of perceptual representations as encoding experience unfolded and the neural mechanisms elicited in the episode offset period to promote the memory formation of episodic sequences. These findings contribute to an emerging understanding of the mechanism by which episodic information is encoded and subsequently retrieved. Current models emphasized the dynamic representational formats through which the brain transforms our experience into a memory trace. Our findings contribute to this important issue by showing that different representational formats are flexibly used during online and offline periods by the brain to support memory formation for episodic sequences.

## Acknowledgements

We thank Bernhard Staresina for helpful discussions in all stages of the study project. This work was supported by the Spanish Ministerio de Ciencia, Innovación y Universidades, which is part of Agencia Estatal de Investigación (AEI), through the project PID2019-111199GB-I00 (Co-funded by European Regional Development Fund. ERDF, a way to build Europe), to L.F. We thank CERCA Programme/Generalitat de Catalunya for institutional support.

